# Bidirectional modulation of reward-guided decision making by dopamine

**DOI:** 10.1101/2024.03.27.586793

**Authors:** Ana Antonia Dias Maile, Theo OJ Gruendler, Adrian G Fischer, Hannah Kurtenbach, Luca F Kaiser, Monja I Froböse, Gerhard Jocham

## Abstract

**Rationale:** The neuromodulator dopamine is known to play a key role in reward-guided decision making, where choice options are often characterized by multiple attributes. Different decision strategies can be used to merge these choice attributes with personal preferences (e.g., risk preferences) and integrate them into a single subjective value. While the influence of dopamine on risk preferences has been investigated, it is unknown whether dopamine is also involved in arbitrating between decision strategies.

**Objective:** In the present study, we investigate the effects of pharmacological dopamine manipulations on arbitrating between different decision strategies in a healthy sample.

**Methods:** 31 healthy male participants performed a reward-guided decision-making task under the influence of the dopamine D_2_/D_3_-receptor antagonist amisulpride (400 mg), the dopamine precursor L-DOPA (100 mg L-DOPA + 25 mg cardidopa), or placebo in a double-blind within-subject design. The effect of dopamine on reward-guided decisions and decision strategies was analyzed using hierarchical implementations of regressions and Bayesian models.

**Results:** Notably, we observed that the dopaminergic interventions shifted the (overall) weighting of option attributes without changing how option attributes are integrated into a subjective value (decision strategy). These effects were bidirectional: Amisulpride *reduced* whereas L-DOPA *increased* the degree to which choices were influenced by both reward magnitude and reward probability. These effects occurred in the absence of changes in statistically optimal behavior.

**Conclusion:** Together, our data provide evidence for a role of dopamine in controlling the influence of value parameters on choice irrespective of decision strategies.

## Introduction

Consider the daily commute to work and the choice of transportation: While biking boosts environmental and financial benefits, it is less comfortable and restricted by weather conditions. In contrast, driving provides flexibility and comfort, yet it has environmental and financial drawbacks. The choice of transportation thus follows from the direct comparison of various choice attributes, demanding the use of decision strategies and attribute weighting (e.g., risk preferences). Two lines of evidence suggest that decision strategies and preferences are controlled by dopamine. First, patients with the neurodegenerative disorder Parkinson’s Disease (PD), which is characterized by a profound dorsal striatal dopamine depletion, exhibit motoric deficits (e.g., tremor and bradykinesia) often accompanied by an increase in risk aversive behavior (Cherkasova et al., 2019; Kobayashi et al., 2019). Conversely, treating the dopamine deficit with the dopamine-precursor levodopa (L-DOPA; Connolly & Lang, 2014) improves the motor symptoms but it can also increase risk seeking behavior and even cause pathological gambling (Evans et al., 2009; Molina et al., 2000; Weintraub et al., 2006).

Second, pharmacological studies in healthy volunteers have shown increased risk seeking after increasing dopamine concentrations by administering L-DOPA (Rutledge et al., 2015). Conversely, similar effects were found when administering the D2/D3-receptor antagonist amisulpride (Burke et al., 2018). Importantly, dopaminergic effects on risk preferences are valence-specific and were exclusively observed in the context of rewards, not losses (Burke et al., 2018; Rutledge et al., 2015). In previous work, choices were usually modelled using Prospect Theory, which posits that, following a non-linear distortion, reward probabilities and magnitudes are multiplicatively combined into an expected value (EV) which is then used to guide decisions (Kahneman & Tversky, 2013; Tversky & Kahneman, 1986). However, an alternative possibility would be that participants do not use such integrated EV at all, but instead rely on simpler heuristics, such as directly comparing attributes and using a weighted combination of attribute differences. Recent evidence suggests that choices of both humans and non-human primates are best characterized by a mixture of such an additive and a multiplicative strategy (Farashahi et al., 2019; Figner & Voelki, 2004; Scholl et al., 2014). These results further suggested that the degree to which either of the two strategies prevails varies as a function of uncertainty about option attributes (Farashahi et al., 2019). Despite the known role of dopamine in both decision making and risk preferences, the effects of dopamine on arbitrating between decision strategies remain untested.

Therefore, in the present study we investigated the role of dopaminergic activity on decision strategies in reward-guided decision making, combining it with computational modeling approaches that take into account both multiplicative and additive strategies for value computation. Additionally, we sought to clarify the dopaminergic effect on risk preferences by including a parameter reflecting the relative balance between reliance on reward probability versus reward magnitude. Healthy participants performed a reward-guided decision-making task under the influence of either the D2/D3-receptor antagonist amisulpride (400 mg), the dopamine precursor L-DOPA (100 mg L-DOPA + 25 mg Carbidopa), or placebo in a double-blind within-subjects design. Notably, there was no significant dopaminergic effect on participants’ decision strategies or risk preferences. Instead, the weighting of choice attributes was shifted. The degree to which participants’ choices were governed by both reward magnitude and reward probability was diminished under amisulpride, but increased under L-DOPA.

## Methods

### Participants

A total of 33 participants took part in the study. Due to the pharmacological manipulation, participants were pre-screened on a separate day to exclude certain pre-existing medical conditions, including neurological or psychiatric disorders. Additionally, an electrocardiogram (ECG) was obtained from each potential participant and assessed by a cardiologist. Only healthy participants without ECG abnormalities were allowed to participate in the study. Due to the menstrual cycle-dependent interaction between gonadal steroids and the dopaminergic system (Becker et al., 1982; Creutz & Kritzer, 2004; Dreher et al., 2007), we only included male participants, as in previous work (Jocham et al., 2011; Jocham, Klein, et al., 2014). Further exclusion criteria were drug abuse, and use of psychoactive drugs or medication in the two weeks before the experiment. Participants were instructed to abstain from alcohol and any other drugs of abuse during the entire course of the study.

Out of the initial 33 participants, two did not fully complete the study due to technical issues leading to the final sample of 31 participants. They were right-handed with normal or corrected-to-normal (*N* = 12) vision. On average, they were *M* = 25.71 years old (age range 21-32, *SD* = 3.20), body weight was *M* = 77.16 kg (weight range 62 – 90kg, *SD* = 7.46). Six participants reported occasional smoking. All were naive to the purpose of the study and gave written informed consent. The present study was approved by the local ethics committee of the Medical Faculty of the Otto-von-Guericke-University Magdeburg (internal reference: 129/13) and is in line with the Helsinki Declaration of 1975. Participants were compensated for their efforts at a fixed rate, plus an additional bonus that depended on their performance during the decision-making task.

### Procedure

The study comprised three pharmacological MEG-sessions in a pseudorandomized order across participants. MEG recordings will be disregarded for the purpose of the present study. Identical procedures were followed on the three sessions, only the drug (or placebo) administered differed. A physician screened participants before each session. In a double-blind cross-over design, participants received a single oral dose of either the dopamine D2/D3-receptor antagonist amisulpride (400mg), the dopamine precursor levodopa (L-DOPA; 100 mg L-DOPA + 25mg cardidopa), or placebo. Because of the different pharmacokinetics of L-DOPA and amisulpride, a dummy administration procedure was used. Participants ingested two pills separated by two hours, of which at least one always contained placebo. In the amisulpride condition, the active substance was contained in the first pill, whereas it was contained in the second pill in the L-DOPA condition. The decision-making task began approximately 1 h after ingestion of the second pill, corresponding to 1 h after L-DOPA administration and 3 h after amisulpride administration in line with the average time for the two drugs to reach peak plasma concentration (Le Bricon et al., 1996; Nutt, 2008). Immediately prior to the reward-guided decision-making task, participants completed Bond & Lader visual analogue scales (BL-VAS; Bond & Lader, 1974) and the trail-making task (TMT, Reitan, 1958) to assess drug effects on mood and visual attention. Heart rate and blood pressure were monitored prior to the first drug (or placebo) administration, prior to the task and after the study, before participants were released by the study physician. Sessions were separated by at least 8 days to ensure complete washout of the drug before the next session (elimination half-life of amisulpride 12 h, L-DOPA with cardidopa 1.5 h).

### Reward-guided decision-making task

Participants completed 500 trials of a reward-guided decision-making task plus 12 additional trials to familiarize themselves with the task beforehand (Figure 1a). On each trial, choices were made between two options defined by two attributes each: a reward magnitude and a probability to obtain this reward. Reward magnitudes were presented as the width of a rectangular horizontal bar and reward probabilities as a percentage written underneath these bars. Therefore, both attributes were explicit and did not have to be learned. Reward outcomes were independent of each other, meaning either of the two options, both, or none of them could be rewarded. Option values were drawn from a reward schedule that was generated before the experiment, as in previous studies (Hunt et al., 2014; Hunt et al., 2013; Jocham, Furlong, et al., 2014; Jocham et al., 2012). This reward schedule was designed such that correlations between factors of interest (in particular between chosen and unchosen and between left and right option value) was minimized. Furthermore, we ensured that reward magnitude and probability were never identical for the two options. To make advantageous choices, participants needed to multiply magnitude and probabilities into an integrative value estimate referred to as EV. On some trials, however, both magnitude and probability of one option were higher than on the alternative option. We refer to these trials as ‘no brainer’ trials. These were limited to occur no more than in 17.8 % of the 500 trials (89 trials).

**Fig. 1.**
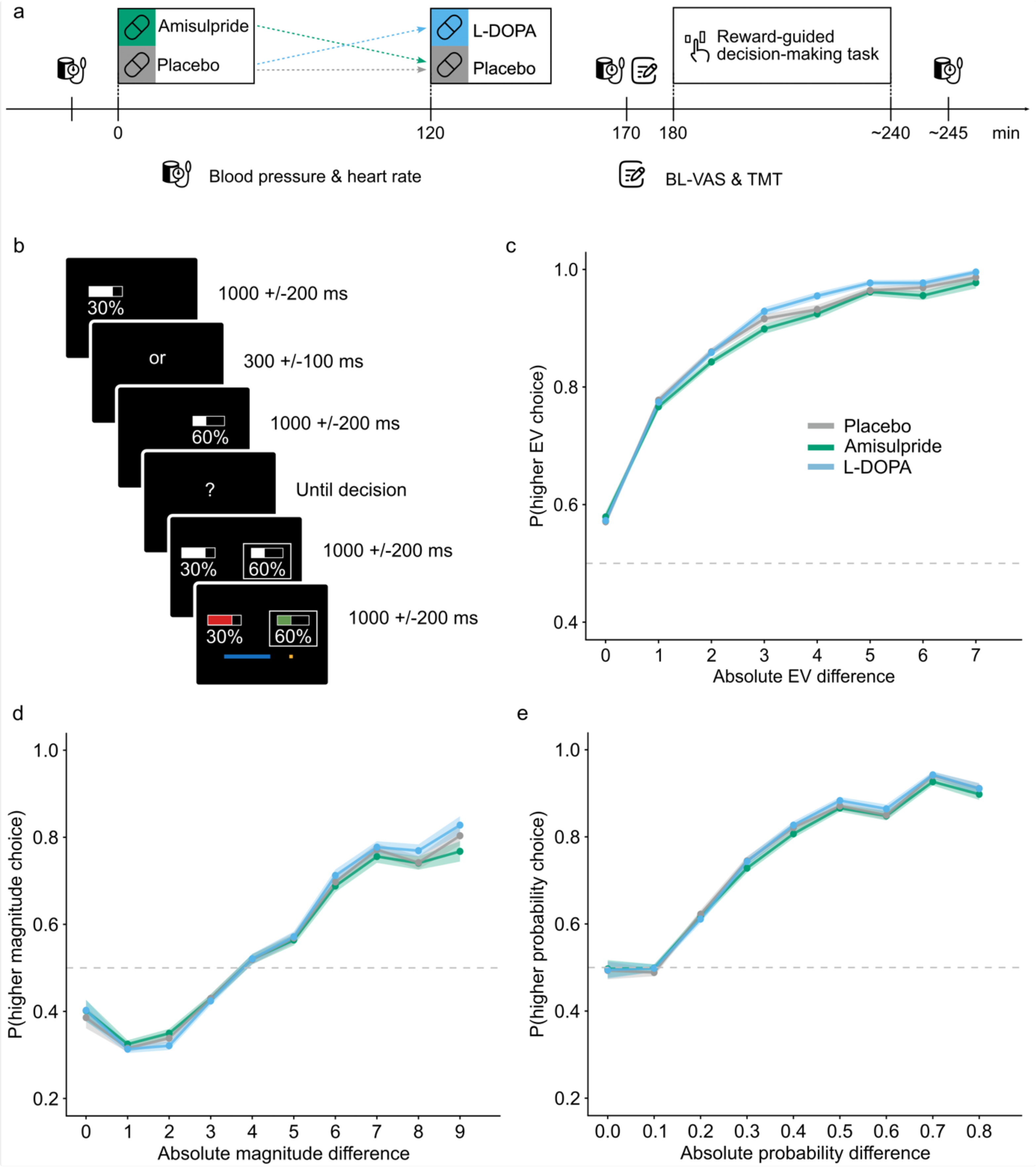
**Overview of study procedure, the experimental task and participants’ behavior**. (a) Study procedures was standardized across all sessions, only the drug (or placebo) administered differed. Due to the different pharmacokinetics of L-DOPA and amisulpride, a dummy administration procedure was used: Participants ingested two pills separated by two hours, of which at least one always contained placebo. Approximately 3 h after the first pill and 1 h after the second pill, participants performed the reward-guided decision-making task. Heart rate and blood pressure were monitored at three different time points. Immediately prior to the task, participants completed Bond & Lader visual analogue scales (BL-VAS) and the trail-making task (TMT). (b) Task schematic and trial timeline. Each choice option was associated with a reward magnitude (width of the horizontal bar) and a reward probability (percentage). The left option was always presented first, followed by a short delay before the presentation of the right option. Participants were instructed to only respond once the question mark appeared. Lastly, they were given feedback on the outcome (reward/no reward) of both the chosen and the unchosen option and reward, if obtained, was added to the progress bar (blue bar at the bottom of the screen). (c) Probability of choosing the choice option with a higher expected value (EV) depending on the absolute EV difference between the two options. Note this is not the mean-centered EV and thus not independent of reward magnitude or probability. (d) Probability of choosing the option with higher reward magnitude depending on the absolute magnitude difference between the two options. (e) Probability of choosing the option with a higher reward probability depending on the absolute probability difference between the two options. In b-d, solid lines represent the average, shaded areas represent SEM across participants.

The two options were presented sequentially such that on every trial, the left option was always presented first, followed by a short delay (200 ms - 400 ms), which was followed by a presentation of the second option on the right side. Immediately after offset of the second option, a question mark appeared prompting participants to indicate their choice. Participants were instructed to refrain from responding while the second option was onscreen and only respond after the question mark had appeared. This design choice was made in order to separate sensorimotor cortical value representations from the execution of movement in the MEG signal. Note that this design precludes a meaningful analysis of reaction times.

Options were selected by button presses with the index finger of the left and right hand, respectively. If an option was rewarded on the current trial, the bar representing the reward magnitude turned green, if it was not rewarded, it turned red. When the chosen option was rewarded, an amount proportional to the reward magnitude of that option was added to the bonus and a blue progress bar displayed at the bottom of the screen indicated this to the participant. To motivate participants, the progress bar displayed earnings towards 2 € and reset itself after reaching this goal. All earned points counted towards the reward. Rewards were rounded to the next higher value in Euro. Even though the outcome of the unchosen option was irrelevant to the bonus, the outcome of both the chosen and the unchosen option was always presented to illustrate independence of outcomes and stimulate risk seeking (i.e., occasional win of low reward probabilities). Outcome presentation was followed by an inter-trial interval (blank screen). The stimuli were presented on a grey (RGB: 60, 60, 60) background with a contrast optimized for the MEG recording chamber (white: 130, 130, 130; green: 46, 139, 60; red: 178, 70, 70; blue: 5, 105, 204).

### Statistical analyses

#### Mixed-effects modeling

Data analyses were conducted using R (R version 4.2.2; R Development Core Team, 2022). All data and codes are made available on OSF (https://osf.io/rnc94/). To evaluate drug effects on decision making, behavioral data was first analyzed using linear mixed modeling, implemented in the lme4 package (version 3.1-3; Bates, Mächler et al., 2015). In contrast to analyses of variance (ANOVAs) and linear regressions, this approach allows taking into account both within- and between-subject variability (Brown, 2021; Harrison et al., 2018; Jaeger, 2008). Since the dependent variable was choice (left vs. right), we used general linear mixed effects models with a binomial distribution. Fixed effects included drug (amisulpride vs. placebo and L-DOPA vs. placebo), the option values (magnitude, probability, mean-centered EV) and a repetition bias (i.e., a tendency to choose the same choice side as in the trial before irrespective of choice attributes). Additionally, the model encompassed the interaction of drug with the option values and repetition bias. To analyze the effects of drug on option values, these were converted into the z-scaled difference scores between the two options. Hence, a drug-induced shift in risk preferences would be evident from a differential (or asymmetric) change of the interaction regression weights for reward magnitude and probability, respectively. To reduce the risk for Type I error and increase power, it is recommended to use the full random effect structure (Barr et al., 2013). However, due to convergence issues, commonly following from an overly complex random effect structure for the data (Bates, Kliegl et al., 2015), we followed the recommendation by Bates, Kliegl et al. (2015) and ran a PCA to inspect the amount of explained variance. Random components that explain 0.00% of variance were not kept in the model. This approach allows systematically maximizing the random effect structure. Hence, linear mixed models included both a random intercept and a random slope for the within-subject factor drug and the option values (magnitude, probability, mean-centered EV). To investigate the task effects irrespective of drug, a separate linear mixed model was set up that included the option values and a repetition bias but disregarded the factor drug. Furthermore, where we observed significant drug effects on control measures (BL-VAS, TMT, heart rate and blood pressure), these effects were added to the task models as fixed effects and we verified that the effects of interest remained unchanged. A further control analysis encompassed adding the session number as fixed effects to control for putative order effects. All *p*-values are based on asymptotic Wald tests. R^2^ is reported for all models using the r.squaredGLMM procedure of the MuMIn package (version 1.48.4; Bartoń, K., 2024).

#### Computational modeling

To analyze the drug effects on the underlying cognitive processes determining participants’ choice behavior, we used a Bayesian hierarchical model (similar to Swart et al., 2017). This method effectively incorporates the within-subject design enabling estimation of the group-level and subject-level parameters. However, due to practicability, we first identified the best fitting model using non-hierarchical models. The first model, the multiplicative model, assumes that reward magnitudes and probabilities are multiplicatively combined (Farashahi et al., 2019):

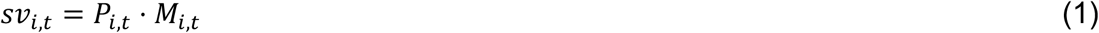

Where *sv_i,t_* is the subjective value, and *M_i,t_* and *P_i,t_* are the reward magnitude and probability, respectively, of option *i* presented on trial *t* (Farashahi et al., 2019).

In contrast, in the additive model, subjective values are computed from a weighted additive combination of reward magnitudes and probabilities:

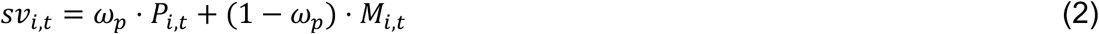

Where ω*_p_* is the relative weight given to probabilities relative to magnitudes. Values of ω*_p_* = 0.5 thus indicate equal influence of probability and magnitude. Values of ω*_p_* > 0.5 indicate that choices are more strongly influenced by probabilities than magnitudes, indicating risk aversion. In contrast, values of ω*_p_* < 0.5 indicate risk seeking (Scholl et al., 2014). For this purpose, magnitudes were scaled to values between 0.10-1.00 aligning them to the value range of reward probabilities.

The third model, the hybrid model assumes that participants use a weighted combination of both the multiplicative and additive strategy:

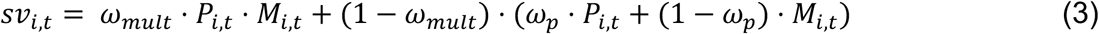

Where the relative allocation between the multiplicative and additive response strategy is governed by the parameter ω*_mult_* (Farashahi et al., 2019; Scholl et al., 2014), which is bound between 0.00 - 1.00, with higher values of ω*_mult_* indicating a dominance of the multiplicative response strategy. In all models, choices were modelled using a softmax choice rule including an inverse temperature *τ* to capture choice stochasticity, which then generates a probability to choose action *A* (in this case, left or right):

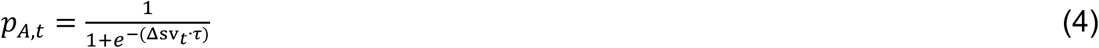

Where *Δsv* is the subjective value difference (left minus right for *A* = left choice).

The linear mixed effect models revealed a (negative) choice repetition bias (see results). Therefore, we extended the best-fitting model by adding a choice bias. Furthermore, to relate to previous work and compare model fits, we used Prospect Theory. This included models where either only reward magnitude (equation 5), only reward probability (equation 6), or both (equation 7) were non-linearly distorted.

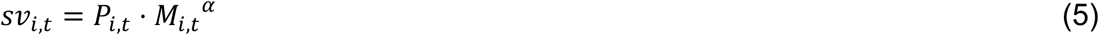

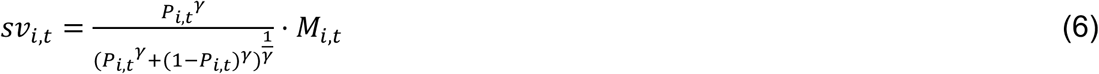

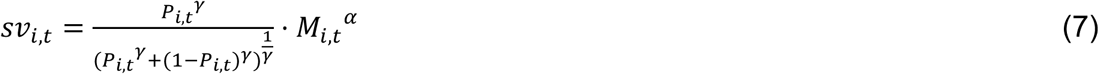

Where *α* and *ψ* are the parameters that describe the non-linear warping of objective into subjective attributes. For this purpose, reward magnitudes were not rescaled and thus ranged between 1-10. Models were implemented using the NLoptr package (version 2.0.3; Johnson, 2008) with bound optimization by quadratic approximation (bobyca; Powell, 2009). The maximum number of function evaluations (*maxeval*) was 10000 and the stopping criterion for relative change (*xtol_rel*) was set to 1.0e-08. Models were run with 100 iterations. The selection of the best-fitting model was based on the Bayes Information Criterion (BIC) for all testing sessions.

Next, the best-fitting model was implemented as a Bayesian hierarchical model to analyze the effects of the dopaminergic drugs. In this approach, group-level parameters (*X*) provide priors for the individual-level parameters (*x*), with *x* ∼ *N*(*X*, *σ*). A half-Cauchy with a scale of 2 defined the hyperpriors for *σ*. With the exception for *X_β_*, the hyperpriors for X were centered around 0: *X_mult,p_* ∼ *N*(0, 2), *X_β_* ∼*N*(2, 3). An inverse logit transform constrained *ω_mult_* and *ω_p_* between 0.00-1.00. The inverse temperature *τ* was positively bounded through an exponential transform. Parameter initials were chosen by training the model on an independent dataset using a similar task (Kurtenbach et al., 2024). We further extended the model by incorporating six parameter shifts (three per drug), enabling analysis of drug-dependent effects on the model parameters (*ω_mult_*, *ω_p_* and τ). These were unconstrained and their hyperpriors were specified by N(0, 3). Initial values of the parameter shifts were set to 0.00. Additionally, we used a similar approach for the best-fitting of the three variants of the Prospect Theory model, enabling a comparison of the effects of the dopaminergic drugs within a flexible attribute weighting model with the traditionally used model (Figure S2). Bayesian hierarchical models were implemented using the RStan package (version 2.21.8; Stan Development Team, 2023), which enables full Bayesian inference with Markov chain Monte Carlo (MCMC) sampling methods. There were four Markov chains, comprising 500 warm-up iterations and 2500 post burn-in iterations per chain (10000 total). The target average acceptance probability (adapt_delta) was set to 0.97. Model convergence was verified using the convergence and diagnostics criteria provided by RStan which include *R̂* < 1.05.

To evaluate the best-fitting model’s ability to capture our data accurately, we conducted posterior predictive checks (Baribault & Colins, 2023). We generated 500 simulated datasets based on the posterior distributions of subject-level parameters obtained from the Bayesian hierarchical model. These simulated datasets were compared and correlated with key behavioral features of the real data, e.g., likelihood of choosing the option with a higher EV, reward magnitude or reward probability.

## Results

### Bidirectional modulation of the reliance of choices on reward information by amisulpride and L-DOPA

In the reward-guided decision-making task, participants were more likely to choose the option with a higher EV, reward magnitude or probability, as evidenced by the descriptive statistics (Figure 1c-e). This is also confirmed by the main effects of task manipulation in the linear mixed models: EV, magnitude and probability, operationalized as a difference score between the two options, have a significant main effect on choice behavior (EV: *β* = 0.90, *SE* = 0.12, *z*(39671) = 7.42, *p* < 0.001; magnitude: *β* = 2.51, *SE* = 0.24, *z*(39671) = 10.31, *p* < 0.001; probability: *β* = 3.79, *SE* = 0.26, *z*(39671) = 14.37, *p* < 0.001; marginal *R^2^_GLMM_* = 0.672; table S1). Accordingly, participants seem to weight magnitudes and probabilities more strongly than the EV as indicated by the numerically higher regression weights. As for the dopaminergic interventions, amisulpride decreased the effect of both reward magnitude and probability on choice (*β* = -0.12, *SE* = 0.05, *z*(39655) = -2.26, *p* = 0.024; and *β* = -0.23, *SE* = 0.07, *z*(39655) = -3.38, *p* = 0.001; for the interaction of amisulpride with magnitude and probability, respectively, marginal *R^2^_GLMM_* = 0.674, Figure 2a-b). In contrast, L-DOPA increased the weighting of reward magnitude and probability on choice (*β* = 0.17, *SE* = 0.06, *z*(39655) = 2.94, *p* = 0.003; and *β* = 0.21, *SE* = 0.07, *z*(39655) = 2.86, *p* = 0.004; for the interaction of L-DOPA with magnitude and probability, respectively, marginal *R^2^_GLMM_* = 0.674, Figure 2c-d). Notably, neither of the dopaminergic drugs had an effect on the influence of EV on choice (*β* = -0.03, *SE* = 0.04, *z*(39655) = -0.60, *p* = 0.550; and *β* = 0.02, *SE* = 0.05, *z*(39655) = 0.35, *p* = 0.728; for the interaction of amisulpride and L-DOPA with EV; marginal *R^2^_GLMM_* = 0.674; table S2). Participants displayed a tendency to alternate between right and left choices from trial to trial, evident from a negative regression weight of the repetition bias (*β* = -0.10, *SE* = 0.04, *z*(39671) = -2.53, *p* = 0.011, marginal *R^2^_GLMM_* = 0.672; table S1). However, this bias was not affected by the dopaminergic interventions (*β* = -0.12, *SE* = 0.07, *z*(39655) = -1.65, *p* = 0.099; and *β* = 0.08, *SE* = 0.08, *z*(39655) = 1.00, *p* = 0.316; for the interaction of amisulpride and L-DOPA with the repetition bias; marginal *R^2^_GLMM_* = 0.674; table S2). Irrespectively, due to the significant impact of the repetition bias on choice behavior, it was taken into account in the following computational modeling. For a complete overview of the linear mixed model results see tables S1 and S2.

**Fig. 2.**
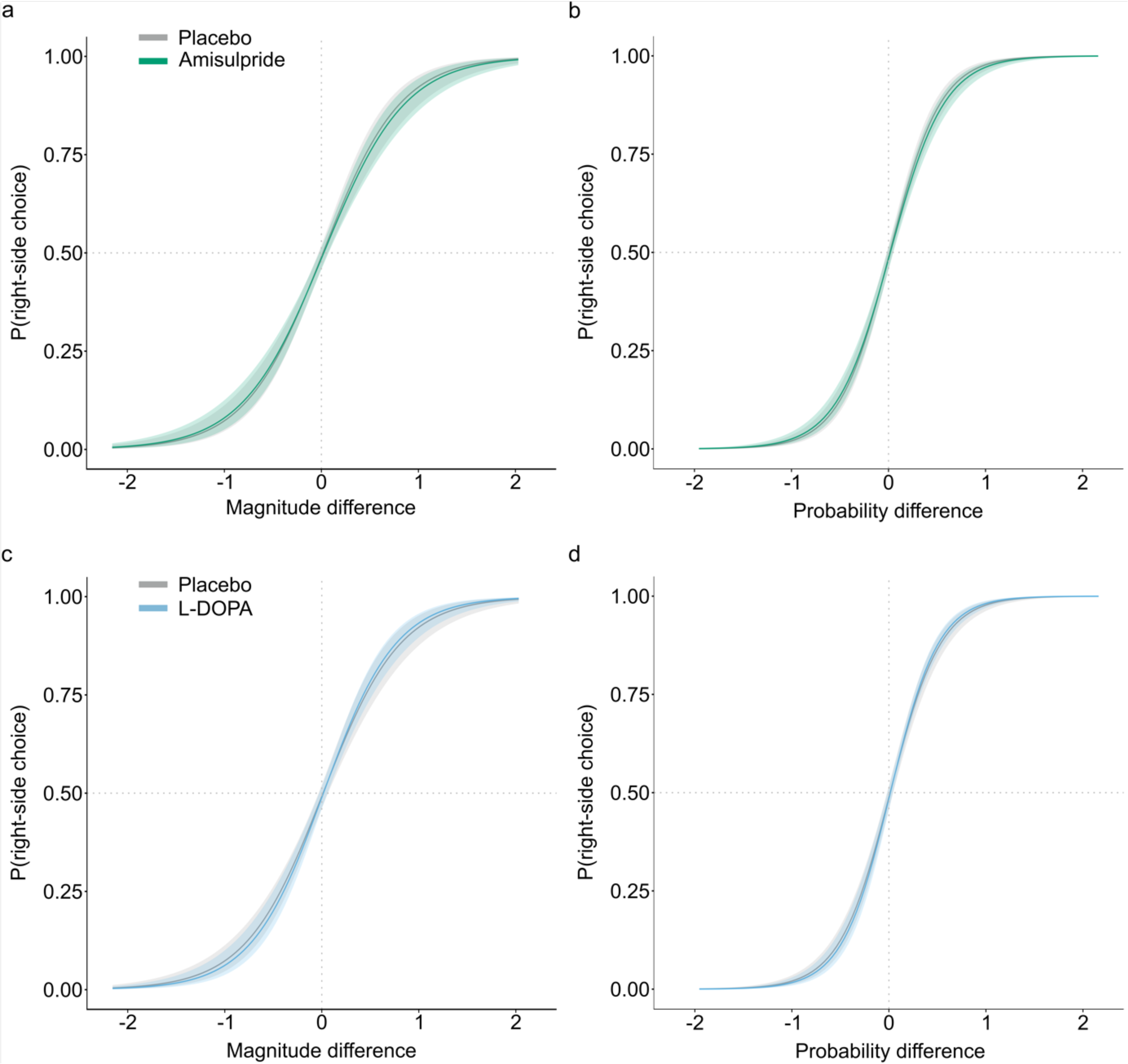
General linear mixed model results. The plots indicate the modelled probability to select the right option as a function of reward magnitude (a, c) or reward probability (b, d) difference between right minus left option (z-scored difference values). Amisulpride significantly decreased the influence of both reward magnitude (a) and reward probability (b) on choice. In contrast, L-DOPA significantly increased the influence of both reward magnitude (c) and reward probability (d) on choice. Solid lines represent the average, shaded areas represent SEM across participants.

We further confirmed that the drug effects were independent of the following potential confounds: All dopaminergic effects were independent of mood, visual attention, heart rate or blood pressure (note that L-DOPA increased participants’ calmness independently of the effects on choice behavior; tables S3-7). When controlling for session order, the reported effects of interest were still significant, however the interaction between amisulpride and reward magnitude was no longer significant (*β* = -0.08, *SE* = 0.06, *z*(39651) = -1.53, *p* = 0.127, marginal *R^2^_GLMM_* = 0.676; tables S8-S9).

### Computational modeling shows no dopaminergic effect on decision strategies

The above results show that amisulpride and L-DOPA changed the degree to which choices were controlled by the individual choice attributes, magnitude and probability. Specifically, amisulpride decreased, and L-DOPA increased the effect of both reward magnitude and probability on choice behavior. Our computational models additionally allow us to further delineate whether these effects arise from an increase in choice stochasticity, as captured by the inverse softmax temperature (*τ*) and/or from a shift in risk preference (*ω_p_*), the relative importance of probability vs magnitude information. Additionally, the computational models allow insights into dopaminergic effects on the selection of decision strategies (*ω_mult_*). The best-fitting model for all testing sessions was the hybrid model without choice bias (BIC *m* = 314.67; Table 1 for model fits across sessions and Table S10 for model fits per session). In this model the parameter *ω_mult_* was estimated as *Mdn* = 0.49 (95%-HDI: 0.37 – 0.61, Figure 3a). This suggests participants used a hybrid response strategy comprising of both a multiplicative and an additive approach instead of a purely multiplicative strategy as traditionally assumed by Prospect Theory. However, this response strategy was not significantly shifted by amisulpride (*Mdn* = 0.02, 95%-HDI: -0.04 – 0.10, Figure 3b) or L-DOPA (*Mdn* = -0.01, 95%-HDI: -0.08 – 0.05, Figure 3B) as the posterior distributions overlap with zero. This indicates no dopaminergic effect on the selection of decision strategies. The weight parameter *ω_p_* reflects the balance between reward probability and magnitude and thus gives insights into participants’ risk preferences. It was estimated to be *Mdn* = 0.82 (95%-HDI: 0.71 – 0.93, Figure 3c). Hence, participants weighted probabilities more strongly than magnitudes causing risk averse behavior. This parameter was not significantly shifted by amisulpride (*Mdn* = 0.04, 95%-HDI: -0.04 – 0.13, Figure 3d) or L-DOPA (*Mdn* = -0.01, 95%-HDI: -0.09 – 0.06, Figure 3d). Therefore, the shifted weighting of choice attributes as observed in the linear mixed models, cannot be attributed to a dopaminergic shift in risk preferences. The inverse softmax temperature *τ* is estimated *Mdn* = 14.56 (95%-HDI: 12.20 – 16.92, Figure 3e). It was not significantly shifted by amisulpride (*Mdn* = -0.46, 95%-HDI: -1.18– 1.02, Figure 3f) or L-DOPA (*Mdn* = 0.62, 95%-HDI: -0.68 – 1.94, Figure 3f). Hence, there is no dopaminergic effect on participants’ choice stochasticity and thus this cannot clarify the dopaminergic shift of choice attributes found in the linear mixed models. Nevertheless, the direction of the drug shifts aligns with the direction of effects found in the linear mixed models: amisulpride decreases the reliance on choice attributes as demonstrated here by the numerically decreasing inverse softmax temperature leading to an increased choice stochasticity. In contrast, L-DOPA increased the weighting of choice attributes aligned with the numerically increased inverse softmax temperature and hence a decreased choice stochasticity. Table S11 depicts the parameter collinearities with all *r_s_* ≤ 0.50. Figure S1 shows the successful posterior predictive checks: the simulated data from the hybrid model reproduce the key features of our data indicated by the highly significant correlation between the real and simulated choice behavior (all *r_s_* ≥ 0.83 and *p* < 0.001). Further, the best-fitting Prospect Theory model included both a probability and magnitude distortion. Model results can be found in the Figure S2. Similar to the hybrid model, there were no significant drug shifts. Nevertheless, although it was not the best fitting model for our data, there was a trend effect for reduced probability distortion under amisulpride (*Mdn* = 0.11, 95%-HDI: -0.01 – 0.24), where 91.1% of the posterior distribution does not overlap with zero.

**Fig. 3.**
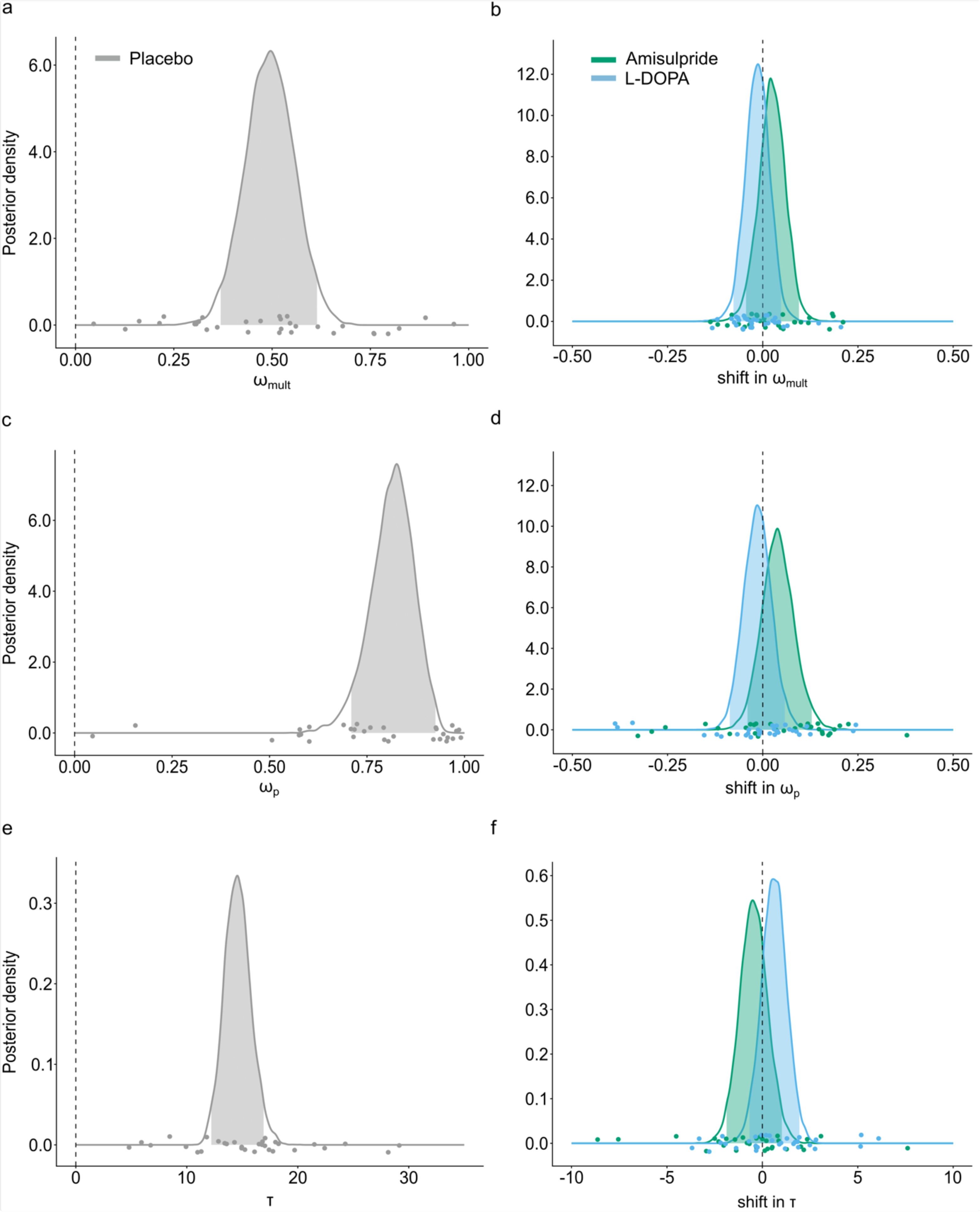
Posterior distributions of the group level estimate of the parameters from the Bayesian hierarchical model. Panels on the left (a, c, e) are the posterior distributions of the three model parameters, the multiplicative weight ω*_mult_*, the relative probability weight ω*_P_* and the softmax inverse temperature τ. Panels on the right (b, d, f) are the posterior distributions of the shift parameters, color-coded for each drug. Shaded areas in the distributions are the 95%-CI. Dots represent single-subject estimates. (a) Participants use a hybrid response strategy comprising of both a multiplicative and an additive decision strategy, as evident from median ω_mult_ = 0.49. (b) This response strategy is not significantly affected by amisulpride or L-DOPA (95%-CI overlaps with zero). (c) Participants choices are guided more by reward probabilities than reward magnitudes, as evident from ω*_P_* higher than 0.5 (median ω*_P_* = 0.82). (d) This relative weighting is not significantly shifted by amisulpride or L-DOPA (95%-CI overlaps with zero). (e) The softmax inverse temperature 1 is not significantly shifted by amisulpride or L-DOPA (f), since the 95%-CI overlaps with zero.

**Table 1.** Model fits for the different computational models. Bayes Information Criterion (BIC) was used to compare model fits. Lower BIC indicates better fit. The hybrid model without a bias is the best fitting model. Values are mean (SE) across all sessions.

|  | BIC |
| --- | --- |
| Hybrid model | 314.67 (10.21) |
| Hybrid model with repetition bias | 319.56 (10.13) |
| Additive model | 336.49 (9.49) |
| Multiplicative model | 400.95 (10.47) |
| Prospect Theory with magnitude distortion | 319.58 (10.38) |
| Prospect theory with probability distortion | 364.93 (10.18) |
| Prospect Theory with magnitude and probability distortion | 317.69 (10.15) |

|  | **BIC** |
| --- | --- |
| **Hybrid model** | 314.67 (10.21) |
| **Hybrid model with repetition bias** | 319.56 (10.13) |
| **Additive model** | 336.49 (9.49) |
| **Multiplicative model** | 400.95 (10.47) |
| **Prospect Theory with magnitude distortion** | 319.58 (10.38) |
| **Prospect theory with probability distortion** | 364.93 (10.18) |
| **Prospect Theory with magnitude and probability distortion** | 317.69 (10.15) |

## Discussion

We investigated the effects of dopaminergic manipulation on reward-guided decision making in a double-blind within-subject design in healthy participants. We found that increasing dopaminergic transmission with L-DOPA increased the degree to which participants’ choices were governed by both option attributes, reward magnitude and probability. Blockade of D_2_-like receptors with amisulpride had the opposite effect, decreasing the effect of both magnitude and probability on choice.

We investigated drug effects using two main approaches. Importantly, we do not intend to suggest that the approaches make independent claims, instead results are complementary, contributing to a more comprehensive understanding. First, linear mixed modeling allowed testing the task parameters that influenced participants’ choices, and how these were affected by drug. We found that the effects of reward magnitude and probability on choice were decreased under amisulpride, and increased under L-DOPA. However, the decreased effect of reward magnitude under amisulpride needs to be interpreted with caution as it was not robust to the control measure session order. In contrast, the weight of the integrated EV was affected by neither of the drugs. Because choosing on the basis of EV represents statistically optimal behavior, this implies that drugs did not affect optimal behavior. We also observed that regression weights for the individual attributes, reward probability and reward magnitude, were higher than those for EV, an issue that we will return to in the next paragraph. Further, in agreement with previous work (Pape & Siegel, 2016; Rogge et al., 2022), we found an alternation bias. That is, above and beyond the effects of value-related parameters, participants displayed a tendency to alternate between right and left choices from trial to trial. However, in contrast to previous studies, this was not modulated by drug (e.g., Rutledge et al., 2009).

Our second approach is based on computational modeling. Traditionally, multi-attribute decision making is captured with models like Prospect Theory (Kahneman & Tversky, 2013; Tversky & Kahneman, 1986). These approaches typically involve applying non-linear weighting functions to reward magnitude and probability prior to multiplying them into an EV. This has recently been challenged by studies showing that behavior of both humans and non-human primates is best characterized by a more flexible mixture of decision strategies (Farashahi et al., 2019; Figner & Voelki, 2004; Scholl et al., 2014), comprised of both EV (multiplicative strategy) and a direct comparison of option attributes (additive strategy). The dominance of either the multiplicative or additive strategy appears to depend on uncertainty where participants have been shown to shift to preferred use of the additive strategy during risky decision making or under heightened uncertainty (Farashahi et al., 2019; Figner & Voelki, 2004). Our modeling results agree with these previous studies. We found that participants’ behavior was best described by a hybrid model incorporating a multiplicative and an additive strategy, which outperformed classic multiplicative models like Prospect Theory. Investigation of the parameter *ω_mult_*, which captures the relative contribution of the two strategies, showed that participants’ subjective values were guided to a similar degree by the multiplicative and additive component. This is in line with the linear mixed modeling regression results, where we also observed that magnitudes and probabilities influenced choice behavior numerically stronger than EV. Further, we reasoned that the dopaminergic change in weighting of the individual attributes, without a concomitant change in the role of EV, should be reflected in choice stochasticity. Therefore, we expected softmax inverse temperature to be decreased under amisulpride, and increased under L-DOPA, corresponding to more and less stochastic choices, respectively. We found no significant effects of amisulpride or L-DOPA on any of the three parameters of the Bayesian hierarchical model.

Nevertheless, numerically, the direction of the drug shifts on the softmax temperature does align with our prediction: Participants’ choice behavior tended to get more stochastic under amisulpride, and less stochastic under L-DOPA. The lack of change in the relative probability weighting parameter *ω_P_* and the multiplicative term *ω_mult_* is again consistent with our regression-based results. A change of *ω_P_* would only be coherent if a drug affected the impact of probability and magnitude in opposite direction - or at the very least if a change in the weight of probability was more/less pronounced than the change in the weight of magnitude. Likewise, a change in *ω_mult_* should have readily been observed in the regression weight for EV.

Dopamine plays a key role in decision making, yet its involvement in arbitrating between multiplicative and additive decision strategies is not known. There is however evidence that dopamine influences risk preferences (Burke et al., 2018; Gross et al., 2021; Ojala et al., 2018; Petzold et al., 2019; Riba et al., 2008; Rigoli et al., 2016; Rutledge et al., 2015; White et al., 2007). For instance, increasing dopamine transmission with L-DOPA has been reported to increase risk seeking behavior (Rigoli et al., 2016; Rutledge et al., 2015). This effect appears to be mediated by participants’ impulsivity and by their sensitivity to reward (Petzold et al., 2019; White et al., 2007). In contrast, D_2_-receptor antagonism has been associated with a decreased risk aversion, as evident from a reduced probability distortion (Burke et al., 2018; Ojala et al., 2018). This is in line with our exploratory analyses on the Prospect Theory model (which was not the best fitting model for our data), where we also found a trend for amisulpride to reduce probability distortion (Figure S2). Our linear mixed modeling effects and the non-significant shift of *ω_P_* in our Bayesian hierarchical model diverge from these previous results. This might be caused by our task design: First, our reward-guided decision-making task consisted exclusively of a gain frame, without a loss frame. Risk preferences, and especially loss aversion, might depend on losing instead of just not receiving a reward. However, previous studies that have also exclusively included a gain frame found dopaminergic effects that were equivalent to studies including both a gain and loss frame (Ojala et al., 2018; Rutledge et al., 2015). Thus, this factor is unlikely to be the reason for our diverging effects. Second, the short offset in our task design between the presentation of the first and second choice option might have modulated the results by imposing a working memory component. The two-state model by Seamans and Yang suggests that dopamine switches prefrontal cortex networks between two states: state 1, dominated by D2-receptor activation, allows exploration of the input space, while state 2, dominated by D1-receptor activation, allows focusing on a limited set of representations leading to enhanced robustness of working memory representations (Seamans et al., 2001; Seamans & Yang, 2004). We cannot rule out that the dopaminergic intervention might have had effects on working memory influencing the differential processing of the first relative to the second option. Third, our sample only included young male participants. This decreases external validity. Yet, studies with a similar pharmacological manipulation and task have found effects in a male-only sample that corresponded well with a mixed sample, suggesting that gender is not a cause for our diverging effects (e.g., Burke et al., 2018; Ojala et al., 2018). Nevertheless, it would be valuable to replicate our findings recruiting a more diverse group of participants. Fourth, similar pharmacological studies have found no effects of (indirect) dopamine agonists or D_2_-receptor antagonists on risk preferences (Acheson & de Wit, 2008; Arrondo et al., 2015; Evers et al., 2017; Hamidovic et al., 2008; Symmonds et al., 2013; Zack & Poulos, 2007).

In conclusion, participants’ behavior was best described by a hybrid model where additive and multiplicative decision strategies jointly contributed to subjective value. Further, our data suggest that dopamine does not affect the relative dominance of either of the two decision strategies. However, the degree to which choices were controlled by individual attributes, was increased by L-DOPA and decreased by amisulpride. Since we did not find evidence for a stronger shift of one option attribute compared to the other, we would argue that this cannot be interpreted as change in risk preferences. Our data thus provide evidence for a role of dopamine in controlling the influence of value parameters on choice, irrespective of decision strategies and risk preferences.

## Acknowledgements

We thank the Department of Neurology at the University Hospital of the Otto-von-Guericke University and the chair of the Department of Neurology, Prof. Dr. Hans-Jochen Heinze for invaluable support. Computational infrastructure and support were provided by the Centre for Information and Media Technology at Heinrich Heine University Düsseldorf.

## Ethical approval

The present study was approved by the local ethics committee of the Medical Faculty of the Otto-von-Guericke-University Magdeburg (internal reference: 129/13) and is in line with the Declaration of Helsinki 1975.

## Consent to participate

Participants gave written informed consent before participation in the study.

## Consent to publish

Not applicable

## Data availability statement

All data and codes are available on OSF (https://osf.io/rnc94/).

## Author contributions

- Study conception and design (TOJG, GJ), data acquisition (TOJG, AGF), data analysis (AADM, HK, LFK, MIF, GJ), interpretation of data for the work (AADM, TOJG, AGF, HK, LFK, MIF, GJ)
- Drafting the article (AADM, MIF, GJ), revising it for intellectual content (AADM, TOJG, AGF, HK, LFK, MIF, GJ)
- Approving the final version of the manuscript (AADM, TOJG, AGF, HK, LFK, MIF, GJ)

