## supplementary materials for "Bidirectional modulation of reward-guided decision making by dopamine"

### Supplementary tables

**Table S1** General linear mixed effects model: task effects. Table depicts the β-weights, *SEM*, *z*-value and *p*-value for the fixed effects. These included main effects for the intercept (side bias), z-scaled differences scores (Δ) between the two options for expected value (EV; constituting of mean-centered values for reward magnitude and reward probability), reward magnitude (M) and probability (P) and a repetition bias. The dependent variable was choice (right-side choice). The marginal $R^2_{GLMM}$ = 0.672. Stars represent significant effects based on $p < 0.05$.

| | ***β-weight*** | ***SEM*** | ***z*-value** | ***p*-value** |
|---|---|---|---|---|
| Intercept | -0.01 | 0.05 | -0.15 | 0.885 |
| ΔEV | 0.90 | 0.12 | 7.42 | < 0.001 * |
| ΔM | 2.51 | 0.24 | 10.31 | < 2e-16 * |
| ΔP | 3.79 | 0.26 | 14.37 | < 2e-16 * |
| Repetition | -0.10 | 0.04 | -2.53 | 0.011 * |

**Table S2** General linear mixed effects model: drug effects. Table depicts the β-weights, *SEM*, *z*-value and *p*-value for the fixed effects. These included main effects for the intercept (side bias), drugs, z-scaled differences scores (Δ) between the two options for expected value (EV; constituting of mean-centered values for reward magnitude and reward probability), reward magnitude (M) and probability (P) and a repetition bias. Additionally, fixed effects included interactions of the drugs with the EV difference, reward magnitude difference, reward probability difference and the repetition bias. The dependent variable was choice (right-side choice). The marginal $R^2_{GLMM}$ = 0.674. Asterisks represent significant effects based on $p < 0.05$.

| | ***β*-weight** | ***SEM*** | ***z*-value** | ***p*-value** |
|---|---|---|---|---|
| Intercept | 0.00 | 0.07 | 0.03 | 0.974 |
| Amisulpride | 0.04 | 0.06 | 0.75 | 0.454 |
| L-DOPA | -0.05 | 0.06 | -0.80 | 0.424 |
| ΔEV | 0.91 | 0.13 | 7.26 | < 0.001 * |
| ΔM | 2.51 | 0.25 | 10.14 | < 2e-16 * |
| ΔP | 3.81 | 0.27 | 14.19 | < 2e-16 * |
| Repetition | -0.10 | 0.05 | -1.83 | 0.068 |
| Amisulpride:ΔEV | -0.03 | 0.04 | -0.60 | 0.550 |
| L-DOPA:ΔEV | 0.02 | 0.05 | 0.35 | 0.728 |
| Amisulpride:ΔM | -0.12 | 0.05 | -2.26 | 0.024 * |
| L-DOPA:ΔM | 0.17 | 0.06 | 2.94 | 0.003 * |
| Amisulpride:ΔP | -0.23 | 0.07 | -3.38 | < 0.001 * |
| L-DOPA:ΔP | 0.21 | 0.07 | 2.86 | 0.004 * |
| Amisulpride:repetition | -0.12 | 0.07 | -1.65 | 0.099 |
| L-DOPA: repetition | 0.08 | 0.08 | 1.00 | 0.316 |

**Table S3** Linear mixed effect model for the pharmacological effects on heart rate. Table depicts the *β-weights*, *SEM*, degrees of freedom, *t*-value and *p*-value for the fixed effects. These included main effects for the intercept, the two drugs (amisulpride and L-DOPA) compared to placebo and the two measuring timepoints (prior to the first drug administration and after the study). Additionally, fixed effects included interactions of the drugs with the measuring timepoint. Asterisks represent significant effects based on $p < 0.05$. The heart rate significantly differs from zero (significant intercept) and decreases from measuring timepoint 1 to 2. There is no significant effect of the drugs on participants' heart rate. The marginal $R^2_{GLMM}$ = 0.257.

| | **β-weight** | ***SEM*** | **df** | ***t*-value** | ***p*-value** |
|---|---|---|---|---|---|
| Intercept | 78.81 | 2.46 | 77.01 | 31.99 | < 2e-16 * |
| Amisulpride | 1.90 | 2.48 | 155.54 | 0.77 | 0.445 |
| L-DOPA | -0.65 | 2.48 | 155.54 | -0.26 | 0.795 |
| Timepoint | -15.39 | 2.51 | 156.11 | -6.14 | < 0.001 * |
| Amisulpride:timepoint | -2.19 | 3.69 | 159.89 | -0.59 | 0.554 |
| L-DOPA:timepoint | 1.13 | 3.69 | 159.89 | 0.31 | 0.759 |

**Table S4** Linear mixed effect model for the pharmacological effects on systolic blood pressure. Table depicts the β-weights, *SEM*, degrees of freedom, *t*-value and *p*-value for the fixed effects. These included main effects for the intercept, the two drugs (amisulpride and L-DOPA) compared to placebo and the two measuring timepoints (prior to the first drug administration and after the study). Additionally, fixed effects included interactions of the drugs with the measuring timepoint. Asterisks represent significant effects based on $p < 0.05$. The systolic blood pressure significantly differs from zero (significant intercept) and there is no significant drug effect. The marginal $R^2_{GLMM} = 0.051$.

| | ***β*-weight** | ***SEM*** | **df** | ***t*-value** | ***p*-value** |
|---|---|---|---|---|---|
| Intercept | 133.85 | 1.96 | 124.42 | 68.30 | < 2e-16 * |
| Amisulpride | -3.26 | 2.39 | 161.43 | -1.36 | 0.175 |
| L-DOPA | -0.49 | 2.39 | 161.43 | -0.21 | 0.838 |
| Timepoint | 3.98 | 2.40 | 162.73 | 1.65 | 0.100 |
| Amisulpride:timepoint | 2.77 | 3.52 | 169.53 | 0.79 | 0.432 |
| L-DOPA:timepoint | -2.02 | 3.52 | 169.53 | -0.58 | 0.566 |

**Table S5** Linear mixed effect model for the pharmacological effects on diastolic blood pressure. Table depicts the β-weights, *SEM*, degrees of freedom, *t*-value and *p*-value for the fixed effects. These included main effects for the intercept, the two drugs (amisulpride and L-DOPA) compared to placebo and the two measuring timepoints (prior to the first drug administration and after the study). Additionally, fixed effects included interactions of the drugs with the measuring timepoint. Asterisks represent significant effects based on $p < 0.05$. The diastolic blood pressure significantly differs from zero (significant intercept) and there is no significant drug effect. The marginal $R^2_{GLMM} = 0.005$.

| | ***β*-weight** | ***SEM*** | **df** | ***t*-value** | ***p*-value** |
|---|---|---|---|---|---|
| Intercept | 83.51 | 1.97 | 107.07 | 42.35 | < 2e-16 * |
| Amisulpride | -1.39 | 2.28 | 159.03 | -0.61 | 0.542 |
| L-DOPA | -2.16 | 2.28 | 159.03 | -0.95 | 0.344 |
| Timepoint | -0.11 | 2.29 | 160.03 | -0.05 | 0.961 |
| Amisulpride:timepoint | 0.59 | 3.37 | 165.81 | 0.18 | 0.861 |
| L-DOPA:timepoint | 1.62 | 3.37 | 165.81 | 0.48 | 0.632 |

**Table S6** Pharmacological effects on mood measured with the BL-VAS and visual attention measured with the TMT. Depicted are the *mean(sd)* of the three factors of the BL-VAS and TMT scores in seconds for each drug condition. Significant deviations from placebo are marked with an asterisks based on $p < 0.05$. L-DOPA significantly increases the factor calmness.

| | **Placebo** | **Amisulpride** | **L-DOPA** |
|---|---|---|---|
| Alertness | 7.41(1.64) | 7.23(1.73) | 7.24(1.68) |
| Contentedness | 7.93(1.57) | 7.92(1.53) | 8.11(1.42) |
| Calmness | 7.51(1.62) | 7.63(2.05) | 8.00(1.63) * |
| TMT | 17.19(5.31) | 17.50(7.25) | 16.40(4.41) |

**Table S7** General linear mixed effects model controlling for the significant effect of L-DOPA on calmness. Table depicts the β-weights, *SEM*, *z*-value and *p*-value for the fixed effects. These included main effects for the intercept (side bias), drugs, z-scaled differences scores (Δ) between the two options for expected value (EV; constituting of mean-centered values for reward magnitude and reward probability), reward magnitude (M) and probability (M), a repetition bias and the calmness scores. Additionally, fixed effects included interactions of the drug with the EV difference, reward magnitude difference, reward probability difference, the repetition bias and the calmness scores. The dependent variable was choice (right-side choice). The marginal $R^2_{GLMM}$ = 0.674. Asterisks represent significant effects based on $p$ < 0.05.

| | ***β*-weight** | ***SEM*** | ***z*-value** | ***p*-value** |
|---|---|---|---|---|
| Intercept | 0.00 | 0.07 | 0.06 | 0.953 |
| Amisulpride | 0.04 | 0.06 | 0.74 | 0.457 |
| L-DOPA | -0.05 | 0.07 | -0.78 | 0.435 |
| ΔEV | 0.91 | 0.13 | 7.26 | < 0.001 * |
| ΔM | 2.51 | 0.25 | 10.15 | < 2e-16 * |
| ΔP | 3.81 | 0.27 | 14.20 | < 2e-16 * |
| Repetition | -0.10 | 0.05 | -1.83 | 0.068 |
| Calmness | 0.02 | 0.05 | 0.42 | 0.671 |
| Amisulpride:ΔEV | -0.03 | 0.04 | -0.60 | 0.551 |
| L-DOPA:ΔEV | 0.02 | 0.05 | 0.35 | 0.730 |
| Amisulpride:ΔM | -0.12 | 0.05 | -2.26 | 0.024 * |
| L-DOPA:ΔM | 0.17 | 0.06 | 2.94 | 0.003 * |
| Amisulpride:ΔP | -0.23 | 0.07 | -3.38 | < 0.001 * |
| L-DOPA:ΔP | 0.21 | 0.07 | 2.86 | 0.004 * |
| Amisulpride:repetition | -0.12 | 0.07 | -1.65 | 0.099 |
| L-DOPA:repetition | 0.08 | 0.08 | 1.00 | 0.318 |
| Amisulpride:calmness | 0.01 | 0.05 | 0.24 | 0.813 |
| L-DOPA:calmness | -0.04 | 0.05 | -0.69 | 0.491 |

**Table S8** General linear mixed effects model: session order effects. Table depicts the β-weights, *SEM*, *z*-value and *p*-value for the fixed effects. These included main effects for the intercept (side bias), drugs, z-scaled differences scores (Δ) between the two options for expected value (EV; constituting of mean-centered values for reward magnitude and reward probability), reward magnitude (M) and probability (P) and a repetition bias. Additionally, fixed effects included interactions of the drugs and the session order with the EV difference, reward magnitude difference, reward probability difference and the repetition bias. The dependent variable was choice (right-side choice). The marginal $R^2_{GLMM}$ = 0.676. Asterisks represent significant effects based on $p < 0.05$.

| | ***β*-weight** | ***SEM*** | ***z*-value** | ***p*-value** |
|---|---|---|---|---|
| Intercept | 0.00 | 0.07 | 0.00 | 0.997 |
| Amisulpride | 0.05 | 0.06 | 0.85 | 0.395 |
| L-DOPA | -0.05 | 0.07 | -0.80 | 0.424 |
| ΔEV | 0.91 | 0.13 | 7.25 | < 0.001 * |
| ΔM | 2.50 | 0.25 | 10.04 | < 2e-16 * |
| ΔP | 3.81 | 0.27 | 14.11 | < 2e-16 * |
| Repetition | -0.10 | 0.05 | -1.78 | 0.076 |
| Amisulpride:ΔEV | -0.02 | 0.04 | -0.48 | 0.634 |
| L-DOPA:ΔEV | 0.01 | 0.05 | 0.28 | 0.783 |
| Amisulpride:ΔM | -0.08 | 0.06 | -1.53 | 0.127 |
| L-DOPA:ΔM | 0.20 | 0.06 | 3.37 | < 0.001 * |
| Amisulpride:ΔP | -0.17 | 0.07 | -2.40 | 0.017 * |
| L-DOPA:ΔP | 0.23 | 0.07 | 3.10 | 0.002 * |
| Amisulpride:repetition | -0.13 | 0.08 | -1.73 | 0.083 |
| L-DOPA:repetition | 0.08 | 0.08 | 1.02 | 0.308 |
| Session:ΔEV | 0.03 | 0.02 | 1.41 | 0.159 |
| Session:ΔM | 0.00 | 0.02 | -0.12 | 0.903 |
| Session:ΔP | 0.22 | 0.03 | 7.67 | < 0.001 * |
| Session: repetition | 0.00 | 0.02 | 0.20 | 0.843 |

**Table S9** General linear mixed effects model: session order effects independent of drug. Table depicts the β-weights, *SEM*, *z*-value and *p*-value for the fixed effects. These included main effects for the intercept (side bias), z-scaled differences scores (Δ) between the two options for expected value (EV; constituting of mean-centered values for reward magnitude and reward probability), reward magnitude (M) and probability (P) and a repetition bias. Additionally, fixed effects included interactions of the session order with the EV difference, reward magnitude difference, reward probability difference and the repetition bias. The dependent variable was choice (right-side choice). The marginal $R^2_{GLMM}$ = 0.675. Asterisks represent significant effects based on $p < 0.05$.

| | ***β-weight*** | ***SEM*** | ***z*-value** | ***p*-value** |
|---|---|---|---|---|
| Intercept | -0.01 | 0.05 | -0.14 | 0.888 |
| ΔEV | 0.91 | 0.12 | 7.40 | < 0.001 * |
| ΔM | 2.53 | 0.25 | 10.28 | < 2e-16 * |
| ΔP | 3.82 | 0.27 | 14.37 | < 2e-16 * |
| Repetition | -0.10 | 0.04 | -2.51 | 0.012 * |
| Session:ΔEV | 0.03 | 0.02 | 1.56 | 0.120 |
| Session:ΔM | 0.01 | 0.02 | 0.30 | 0.762 |
| Session:ΔP | 0.24 | 0.03 | 8.26 | < 2e-16 * |
| Session:repetition | 0.01 | 0.02 | 0.30 | 0.765 |

**Table S10** Model fits for the different computational models per session. Bayes Information Criterion (BIC) was used to compare model fits. Lower BIC indicates better fit. The hybrid model without a bias is the best fitting model in each session. Values present *mean (SE*).

| Session 1 | BIC |
|---|---|
| **Hybrid model** | 341.80 (19.44) |
| **Hybrid model with repetition bias** | 346.23 (19.08) |
| **Additive model** | 362.29 (17.78) |
| **Multiplicate model** | 403.18 (17.05) |
| **Prospect Theory with magnitude distortion** | 344.60 (19.22) |
| **Prospect Theory with probability distortion** | 375.63 (17.63) |
| **Prospect theory with magnitude and probability distortion** | 344.20 (19.34) |
| **Session 2** | **BIC** |
| **Hybrid model** | 298.24 (16.91) |
| **Hybrid model with repetition bias** | 303.31 (16.79) |
| **Additive model** | 319.37 (16.35) |
| **Multiplicate model** | 404.03 (19.80) |
| **Prospect Theory with magnitude distortion** | 304.28 (18.11) |
| **Prospect Theory with probability distortion** | 363.80 (18.37) |
| **Prospect theory with magnitude and probability distortion** | 300.40 (16.91) |
| **Session 3** | **BIC** |
| **Hybrid model** | 303.98 (16.06) |
| **Hybrid model with repetition bias** | 309.14 (16.17) |
| **Additive model** | 327.79 (14.51) |
| **Multiplicate model** | 395.64 (18.02) |
| **Prospect Theory with magnitude distortion** | 309.86 (16.21) |
| **Prospect Theory with probability distortion** | 355.35 (17.25) |
| **Prospect theory with magnitude and probability distortion** | 308.46 (15.85) |

**Table S11** Parameter collinearity. Correlation between the model parameters based on the subject-level posterior distributions.

| | $\omega_{mult}$ | $\omega_p$ | $\tau$ |
|---|---|---|---|
| $\omega_{mult}$ | 1.00 | -0.05 | 0.50 |
| $\omega_p$ | -0.05 | 1.00 | -0.06 |
| $\tau$ | 0.50 | -0.06 | 1.00 |

### Supplementary figures

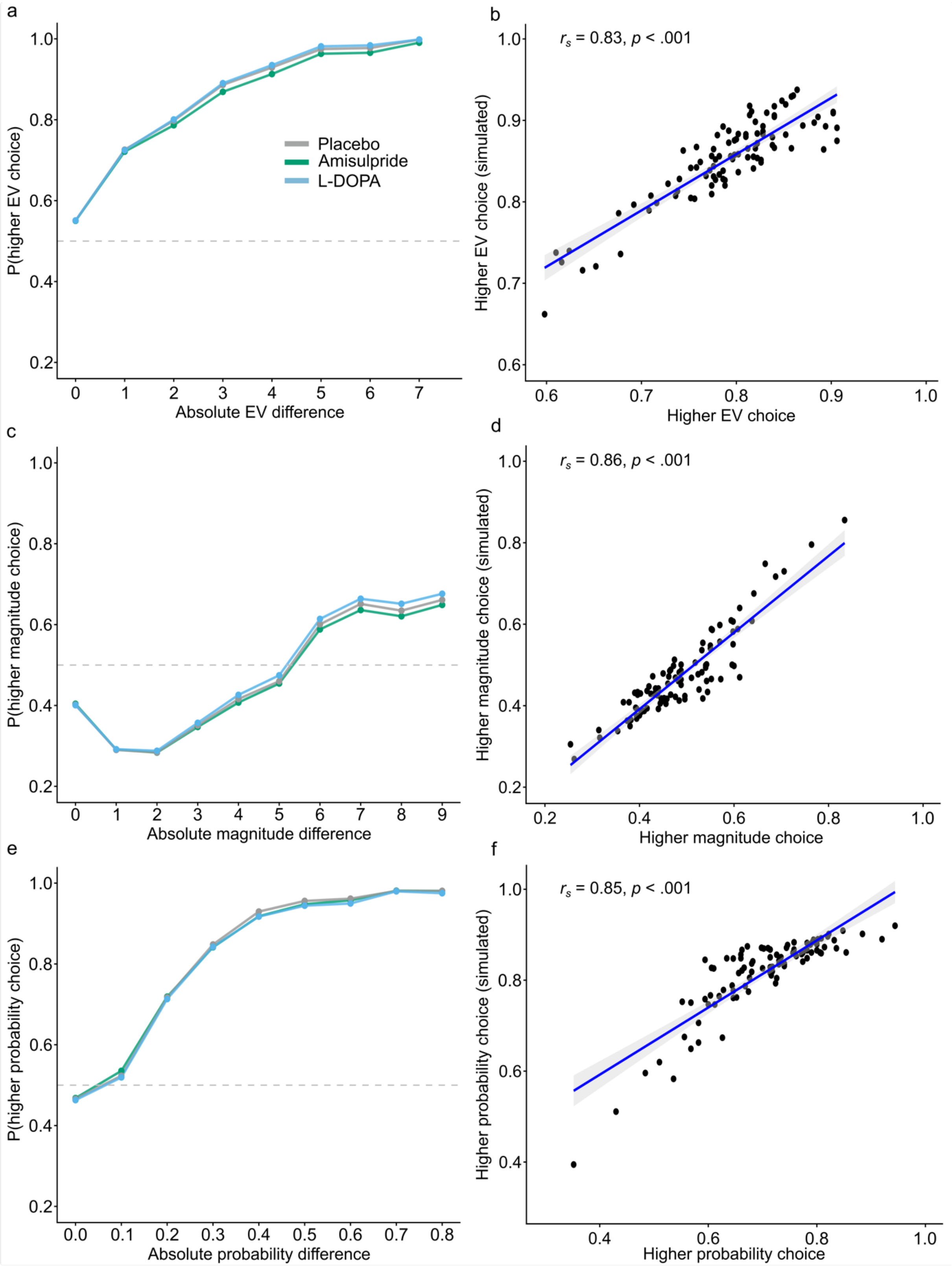

**Fig. S1** Posterior predictive checks. To evaluate the hybrid model's ability to capture the data accurately, we generated 500 simulated datasets based on the posterior distributions of subject-level parameters from the Bayesian hierarchical model. The simulated data is plotted (a, c, e) and correlated (b, d, f) with key features of the real data. In a, c and e drug conditions are color coded. Shaded areas around the graphs depict the *SEM***.** (a), Probability of choosing the choice option with a higher EV depending on the absolute EV difference between choice options. (b), Correlation between real and simulated choices with higher EVs (c), Probability of choosing the choice option with a higher reward magnitude depending on the absolute magnitude difference between choice options. (d), Correlation between real and simulated choices with higher reward magnitudes (e), Probability of choosing the choice option with a higher reward probability depending on the absolute probability difference between choice options. (f), Correlation between real and simulated choices with higher reward probabilities.

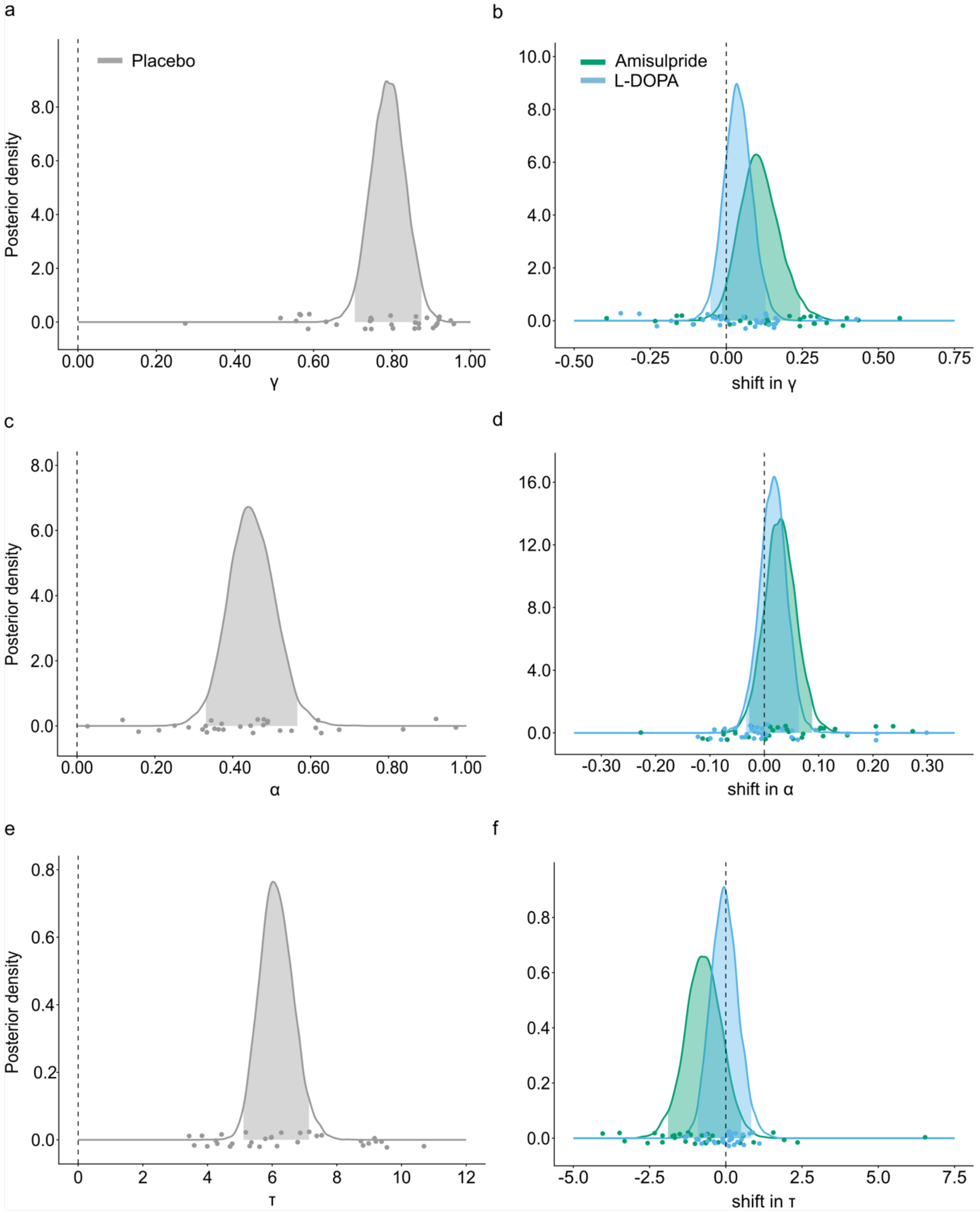

**Fig. S2** Posterior distributions of the group level estimate of the parameters from the Bayesian hierarchical Prospect Theory model. Panels on the left (a, c, e) are the posterior distributions of the three model parameters, the probability distortion γ, the magnitude distortion α and the softmax inverse temperature τ. Panels on the right (b, d, f) are the posterior distributions of the shift parameters, color-coded for each drug. Shaded areas in the distributions are the 95%-CI. Dots represent single-subject estimates. (a) As shown by median γ = 0.79 (95%- HDI: 0.71 – 0.87), participants distort reward probabilities. (b), This probability distortion is not significantly shifted by amisulpride (*Mdn* = 0.11, 95%- HDI: -0.01 – 0.24) or L-DOPA (*Mdn* = 0.04, 95%- HDI: -0.05 – 0.13) since the 95%-CI overlap with zero. Nevertheless, there is a trend effect where amisulpride increases γ since 91,1% of the posterior distribution does not overlap with zero. The increase in γ leads to a reduced probability distortion under amisulpride (c), Reward magnitudes are distorted as shown by median α = 0.45 (95%- HDI: 0.33 – 0.57). (d), This magnitude distortion is not significantly shifted by amisulpride (*Mdn* = 0.03, 95%- HDI: -0.03 – 0.09) or L-DOPA (*Mdn* = 0.02, 95%- HDI: -0.03 – 0.06) since the 95%-CI overlap with zero. (e), The median of the inverse softmax temperature $\tau$ is estimated *Mdn* = 6.12 (95%- HDI: 5.11 – 7.14). (f), The inverse softmax temperature is not significantly shifted by amisulpride (*Mdn* = -0.73, 95%- HDI: -1.90 – 0.50) or L-DOPA (*Mdn* = -0.05, 95%- HDI: -0.92 – 0.84) since the 95%-CI overlap with zero.
